# Multimodal single-cell analyses reveal distinct fusion-regulated transcriptional programs in Ewing sarcoma

**DOI:** 10.1101/2025.06.18.660457

**Authors:** Olivia G. Waltner, April A. Apfelbaum, Emma D. Wrenn, Shruti S. Bhise, Sami B. Kanaan, Rula Green Gladden, Mark A. Mendoza, Roger Volden, Zev Kronenberg, Anand Patel, Michael Dyer, Jay F. Sarthy, Elizabeth R. Lawlor, Scott N. Furlan

## Abstract

Ewing sarcoma (EwS) is a fusion-driven malignancy, peaking in adolescence. Although EwS tumors are driven uniquely by EWS::FLI1 and related fusions, patient outcomes vary greatly. If and how tumor plasticity of EWS::FLI1-regulated transcriptional signatures contribute to disease progression is not known. To address this, we utilized a single-cell co-assay of RNA and chromatin accessibility (ATAC) sequencing to identify gene regulatory networks in EwS. By comprehensively characterizing regulatory elements across cell lines, we identified multiple unique modules of gene regulation. Differential usage and prevalence of these modules was evident across cell lines, associated with distinct epigenetic and transcriptomic signatures, and in specific cases, modifiable by exogenous TGF-β. When we examined primary EwS patient tumors, we observed these same regulatory modules were variably enriched both across and within tumors, highlighting the existence of intratumoral heterogeneity in gene regulatory networks. Our findings demonstrate that multiple, co-existing transcriptional programs shape the phenotypic diversity of EwS and suggest that the balance between these networks may have important implications for clinical outcomes and targeted therapy development.

**Summary:** Multimodal transcriptional analysis reveal how Ewing sarcoma tumors use distinct gene programs, including one linked to TGF-β, to drive cancer behavior and progression.

## Introduction

Ewing sarcoma (EwS) is an aggressive, fusion-driven bone and soft tissue tumor encountered most in adolescents and young adults. Over a third of patients, including nearly all with metastatic or recurrent disease, do not survive^1,2^. Clinical presentation of EwS is heterogeneous, with 80% of localized cases presenting in the bone, and 20% arising in extraosseous locations. Histologically, EwS tumors are highly undifferentiated, exhibiting phenotypic features of both mesenchymal and neural crest progenitors. Although the cell of origin is controversial, EwS tumors are presumed to be derived from these lineages^3–6^. Chromosomal translocations that create fusions between the N-terminal domain of *EWSR1* and the DNA-binding domain of ETS transcription factor (TF) family members are the primary oncogenic lesion in EwS^1^. The resulting EWS::ETS chimeric oncoprotein reprograms gene regulatory networks through diverse mechanisms, including modifying chromatin state through its function as a pioneer factor at GGAA microsatellite repeats, creating *de novo* enhancers^7,8^. Apart from their fusions, EwS tumors are relatively bereft of mutations^1,9^. Neither additional genetic alterations, nor differences in fusion breakpoint and binding partner heterogeneity^10^ adequately explain the diverse clinical outcomes that are observed in patients who receive identical standard-of-care therapies. Advances in transcriptomic and epigenomic analyses have in recent years demonstrated the profound contributions of tumor cell plasticity and cell state heterogeneity to tumor metastasis, relapse, and therapy resistance^11,12^. EwS tumors demonstrate marked inter- and intra-tumoral heterogeneity in their epigenetic states^13,14^ and, as such, these differences may be key determinants of disease progression and clinical outcome.

Recent studies by our group and others have used single cell genomics to study cellular heterogeneity within EwS tumors^14–23^. These studies reveal that EwS tumors coopt distinct developmental programs to promote proliferation or migratory behavior^15,16,18,20,21,24,25^. In addition, accruing evidence shows that cooperativity between TFs, EWS::FLI1, and diverse chromatin states may underlie these various oncogenic programs^20,26–34^. We hypothesized that joint single-cell transcriptional and chromatin profiling technology would reveal new insight into the cell-intrinsic epigenetic and transcriptional heterogeneity of EwS cell states. Using this approach, we have identified a set of diverse gene regulatory networks that are differentially influenced by the EWS::FLI1 fusion and TGF-β in the tumor microenvironment. These distinctive networks vary both across and within tumors from EwS patients providing evidence that a diversity of gene regulatory programs may contribute to the heterogeneity of this disease.

## Results

### Multiomic analyses reveal distinct chromatin accessibility states across EwS cell lines

Cell lines are one of few available models used to interrogate EwS biology given the rarity of the disease and the lack of appropriately faithful animal models^35^. One drawback of cell line use is potential loss of intra-tumoral heterogeneity compared to the original tumor due to serial passaging. We reasoned that profiling an array of EwS cell lines would capture the diversity of gene regulatory networks that may underlie the clinical and transcriptional heterogeneity of this disease. Because of the well-described direct influence of the EwS fusion on chromatin remodeling ^34^, we utilized an assay capable of pairing single-cell RNA transcript counts with a chromatin assay for transposase-accessible chromatin (ATAC) (herein referred to as ‘multiomic’ sequencing) on 9 EwS and 5 non-EwS cell lines (Fig.1A, Fig.S1A). We recovered 30,105 profiles after filtering for poor quality cells (Fig.1A, Fig.S1A, Fig.S1B). Multiomic single-cell profiles self-assembled after dimensionality reduction according to cell line identity (Fig.1A).

**Fig. 1.**
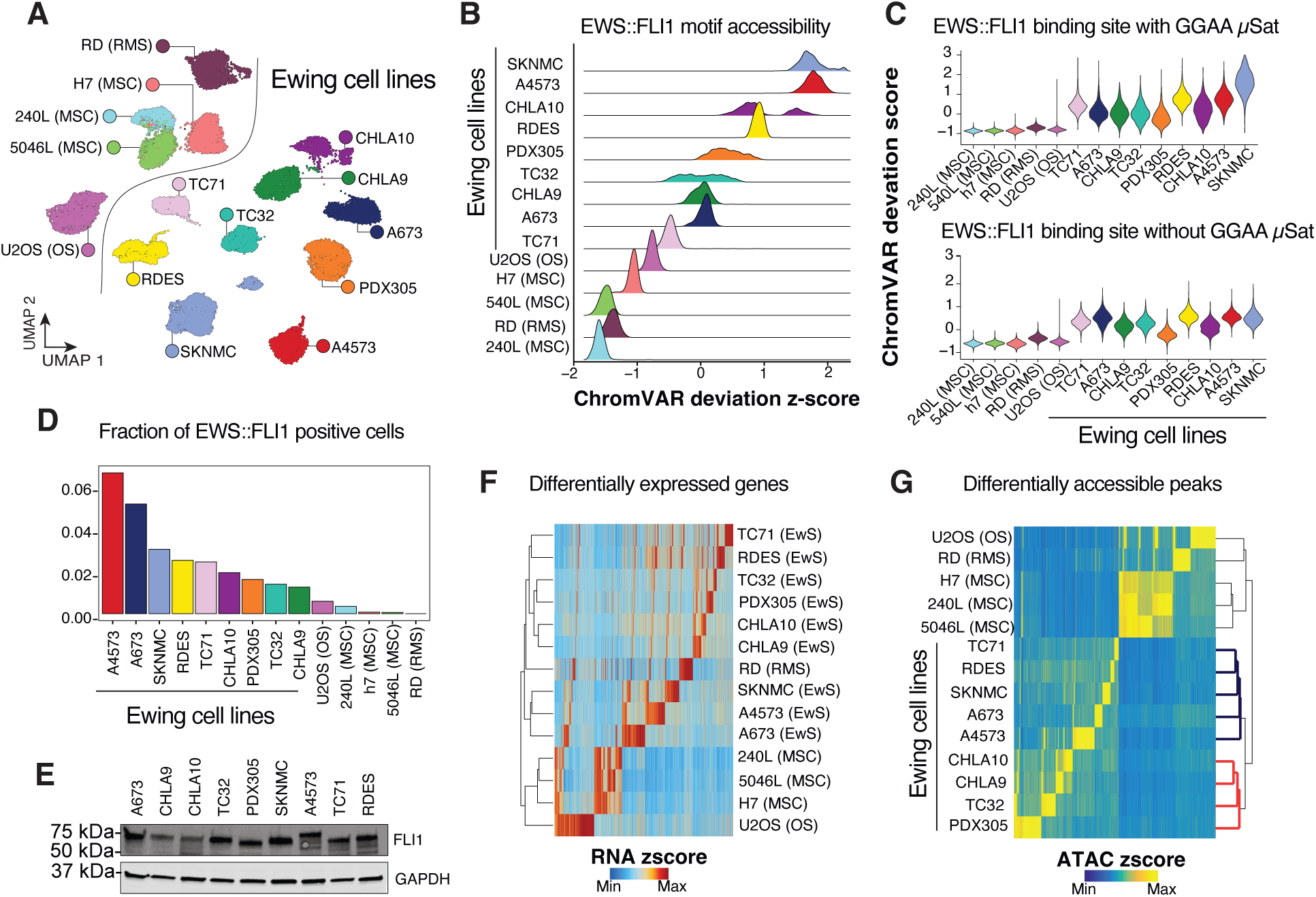
Multiomic analyses reveal distinct signatures between non-EwS and EwS cells. **(A)** UMAP embedding of single cell RNA sequencing and ATAC sequencing of EwS and non EwS cells. **(B)** ChromVAR deviation scores of EWS::FLI1 motif enrichment across EwS and non-EwS cells **(C)** ChromVAR deviation scores of EWS::FLI1 binding site enrichment, separated by regions with or without GGAA microsatellites (μSAT; GGAA repeat ≥ 4), grouped by cell line. **(D)** Fraction of cells in each cell line with detection of EWS::FLI fusion transcript. **(E)** Western blot of FLI1 across EwS cell lines. GAPDH = loading control. **(F)** Differentially expressed genes (FDR ≤ 0.01 & Log2FC ≥ 1.25) across EwS/non-EwS landscape. **(G)** Differentially accessible peaks (FDR ≤ 0.01 & Log2FC ≥ 1.25) of EwS and non EwS cells with row dendrogram showing phenotypic clusters. Row dendrogram shows EwS cell clusters highlighted in bold and red.

First, to identify the extent of transcriptional heterogeneity of validated direct EWS::FLI1 target genes^19^ (EwS TGs) in individual cells, we calculated expression of EwS TGs at the single-cell level, subtracted by the aggregated expression of control gene sets^36^. As expected, RNA expression of these EwS TGs was higher in EwS compared to non-EwS cell lines (Fig. S1C). As these published EwS TGs were experimentally derived from the A673 cell line^19^, it was unsurprising that A673 exhibited the highest average expression of these genes. To interrogate EwS TGs using chromatin-accessibility data, we generated gene activity scores^37^ which model accessibility of regulatory elements in the vicinity of a given gene. These scores confirmed that chromatin accessibility of EwS TGs was higher in EwS cell lines compared to non-EwS cells (Fig. S1D). To compare EWS::FLI1 motif enrichment across cells, we calculated single-cell ChromVAR deviation scores^37^ of GGAA-microsatellites (four or more consecutive GGAA repeats; abbrev. ‘GGAA- µSat’) in accessible regions. As anticipated, this analysis revealed low enrichment in non-EwS cell lines. Unexpectedly, stark differences in EWS::FLI1 motif enrichment were observed across EwS cell lines (Fig. 1B). In some cell lines accessibility of GGAA- µSats was similar across single cells (e.g. TC71, RDES) while others showed a broader range of accessibility (e.g. SKNMC, TC32, PDX305). CHLA10 cells showed a clear bimodal distribution of EWS::FLI1 motif accessibility (Fig. 1B). This hints that within these cell lines, heterogeneity exists among individual cells with respect to accessibility of GGAA- µSats. Next, we collated a list of publicly available EWS::FLI1 fusion binding sites^38^ with or with a GGAA µSat present. We then assessed chromatin accessibility at these published EWS::FLI1 binding sites across all cell lines and discovered that accessibility at these loci varied in a cell-line-specific manner (Fig. 1C & Fig. S1D).

Given the heterogeneity in the magnitude of EWS::FLI1 motif enrichment across EwS cell lines, we hypothesized that EwS cell lines differ in the proportion of cells expressing high levels of the EWS::FLI1 fusion transcript. It has been reported that lower levels of EWS::FLI1 gene expression and activity induce mesenchymal phenotypes in EwS cell lines^39^. To address this, we designed an approach to enrich EWS::FLI1 fusion transcript from the RNA components of our multiomic capture using biotinylated hybridization probes that tiled the EWSR1 and FLI1 gene loci. We then performed long-read sequencing on the product and counted fusion reads using novel fusion detection software designed specifically for long-reads (see Methods for details). As expected, EWS::FLI1 fusion transcripts were detected in EWS cells (Fig.1D). Scant non-ES cells showed low levels of the fusion, likely from PCR artifacts of low-complexity products. Overall, however, we observed that EwS cell lines exhibit a high degree of variability in detectible EWS::FLI1 fusion transcript (Fig. 1D & Fig. S1E), possibly reflecting a difference in expression levels among individual cells. Generally, the proportion of cells with detectible transcript correlated with values obtained using single-cell qPCR to assess fusion expression^15^. These findings are generally congruent with motif enrichment in that high fusion-expressing cell lines (e.g. A4573, SKNMC, and RDES) exhibit high EWS::FLI1 motif enrichment while cell lines bearing lower levels of fusion transcript and FLI1 product (e.g. PDX305, TC32, CHLA9) exhibit lower EWS::FLI1 chromatin signature (Fig.1B & Fig. 1D-E). By contrast, other cell lines displayed inverse trends of fusion transcript and product to chromatin enrichment (e.g. A673 and TC71).

To profile the global chromatin and transcriptional heterogeneity using an unbiased approach, we calculated top differentially expressed (DE) genes and differentially accessible (DA) regions across EwS and non EwS cells. As with the EwS TG-focused analysis, this approach showed that top DE/DA features readily distinguished EwS cells from non-EwS cells (Fig. 1F-G). We observed self-aggregation of EwS cells into distinct clusters. Specifically, in terms of top DA regions, CHLA9, CHLA10, TC32, and PDX305 cells were more similar to each other than other EwS cell lines (Fig. 1G). Altogether, these data show that EwS cell lines exhibit heterogeneous gene-regulatory profiles that segregate into groups that display distinct trends in accessibility, potentially reflecting unique underlying gene regulatory networks that are active in discrete cell contexts.

### EwS cells utilize gene regulatory programs that drive inter-tumoral heterogeneity

Next, to evaluate how cis-regulatory gene networks differ across EwS cells, we correlated transcript expression to regions of accessibility using the joint measurements of transcript count and accessibility in each cell (so-called “peak-to-gene linkages”)^37^. Hierarchical clustering of these cis-regulatory networks with a k parameter equal to 3 clearly separated CHLA9, TC32, and PDX305 into one distinct network module and SKNMC, A4573, RDES, and TC71 into another (Fig. 2A), recapitulating what we observed using top DE/DA features (Fig. 1F-G). We selected k = 3 to extend the two-cluster structure observed in Fig. 1F–G, reasoning that A673 represented a clear outlier whose distinct regulatory program would be obscured at lower k. Cell subsets of CHLA10 separated into either cluster 2 or 3 (Fig. 2A) but more cells had cumulative expression of module 2 genes (Fig. 2B). We then assigned the 9 cell lines into three groups based on their dominant module signature (Fig. 2A-B), resulting in assignment of A673 cells as group 1, CHLA9, CHLA10, TC32, and PDX305 cells as group 2, and SKNMC, A4573, TC71, and RDES as group 3 (Fig. 2A). Enrichment analysis using Gene Ontology and Hallmark terms confirmed that the genes associated with these modules have functional associations with distinct cell behaviors (Fig. 2C). While there was some degree of overlap, module 2 showed a relative enrichment in GO terms related to mesenchymal identity, and modules 1 and 3 had enriched neuronal identity markers. Genes involved in oxidative phosphorylation and targets of mTOR were uniquely enriched in module 1, possibly due to the presence of a BRAF^V2600E^ mutation in A673 cells given the reported effects of this mutation on the mTOR pathway^40–42^.

**Fig. 2.**
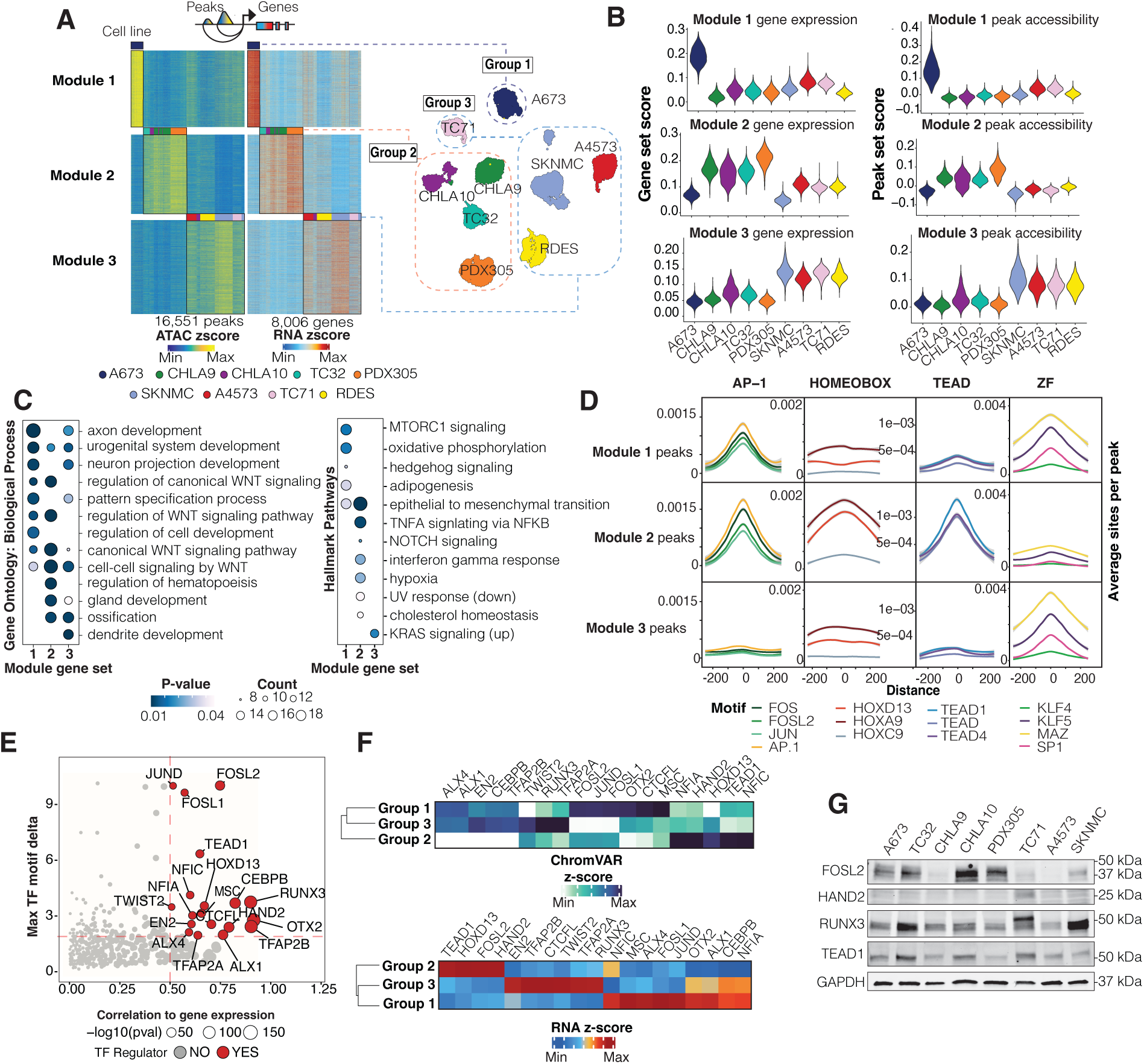
EwS cells utilize distinct gene regulatory modules. **(A)** Heatmap of unsupervised kmean clusters of peak-to-gene links (PGLs) across 9 EwS cell lines. Left- peaks, right-genes, top- colors denote cell lines. Right of heatmap-UMAP of cell lines labeled by subgroup. Group 1 labelled with navy/dashed lines, group 2 in orange and group 3 in light blue. (B) Violin plot of modules 1-3 gene expression and peak accessibility across EwS cell lines. (C) Universal enrichment analysis of module gene (top 300 filtered on peak-to-gene correlation) sets using both MSigDB Hallmark pathways and Gene Ontologies: Biological Process. The top 7 terms filtered by lowest p-value (those greater than 0.05 are not plotted) and count are shown. More significant terms are darker blue. The number of genes in each module that overlap with a gene set term is indicated by the count metric—represented as dot size. (D) HOMER de novo motif enrichment for each module peak set. Results depict the top significantly enriched motifs in each module peak set, and enrichments in other modules. (E) TFs with high expression and motif enrichment in entire single-cell data set of EwS cell lines. Statistically significant TFs correlation to gene expression > 0.5 and a maximum inter EwS subgroup difference in motif deviation zscore (denoted max TF motif delta) > 1.9 highlighted in red. (F) ChromVAR motif deviation scores and gene expression of highly enriched TFs, parsed by EwS subgroup. (G) Western blot of selected TFs across EwS cell lines. GAPDH = loading control.

Next, we investigated if gene modules might harbor distinct TF binding sites. We performed differential motif enrichment using HOMER^43^ and found a strikingly well-delineated, and module-specific enrichment of specific TF families across the cis-regulatory networks (Fig. 2D). The AP-1 family of motifs (FOS, FOSL2, JUN, AP1), which has been suggested to form ternary complexes with EWS::FLI1^44^, enriched in modules 1 and 2, but not in module 3 (Fig. 2D). Zinc Finger (ZF) TFs (KLF4, KLF5, MAZ, SP1) were most enriched in modules 1 and 3 with limited evidence for activity in regulation of module 2 (Fig. 2D). Module 2 peaks uniquely exhibited motif enrichment for developmental regulators from the Homeobox family of TFs (HOXD13, HOXA9) and downstream transcriptional regulators of the Hippo and Wnt signal transduction pathways (TEAD1, TEAD4). To corroborate these findings, we performed an analysis which correlates, at the single-cell level, TF binding motifs in accessible regions with gene expression (Fig. 2E). This showed that TEAD1, HOXD13, and AP-1 TFs were highly enriched in group 2 cells (Fig. 2E-F). Group 3 lacked motif enrichment for AP-1 TFs but showed robust enrichment in ALX1/4, RUNX3, CEBPB and EN2 TFs (Fig. 2E-F). We assessed protein levels of selected TF regulators by Western blots to evaluate their heterogeneity across the cell lines. The protein abundance of these TFs exhibited substantial variability among the cells tested (Fig. 2G). Together, these multimodal analyses of single cells uncovered at least two dominant gene regulatory circuits. One of these (module 2) displays mesenchymal/EMT-like functionality that is likely to be mediated by high activity of TEAD, AP-1, and HOXD13 TFs. The other (module 3) has enriched neural program functionality most likely mediated by ZF TFs and neural and neural crest development TFs such as RUNX3 and HAND2.

### EwS gene regulatory modules are differentially regulated by EWS::FLI1

We next sought to address whether the identified gene regulatory modules are directly or indirectly modulated by EWS::FLI1. We first measured EWS::FLI1, EWS::ERG (the second most common fusion in EwS tumors), and wild-type FLI1 motif enrichment and found all to be enriched in all three modules (Fig. 3A) suggesting that EWS::FLI1 is likely to regulate expression of linked genes across all cells irrespective of the dominant module. However, by annotating these sites using published EWS::FLI1 binding sites (*39*) and segregating into those with or without a GGAA microsatellite (GGAA- µSat), we observed enrichment of GGAA- µSats only in module 3 peaks (Fig. 3B). Thus, this analysis suggests that EWS::FLI1 binding is likely to directly influence gene regulatory circuits and its impact on transcriptional state depends on GGAA patterning, as has previously been suggested^7^.

**Fig. 3.**
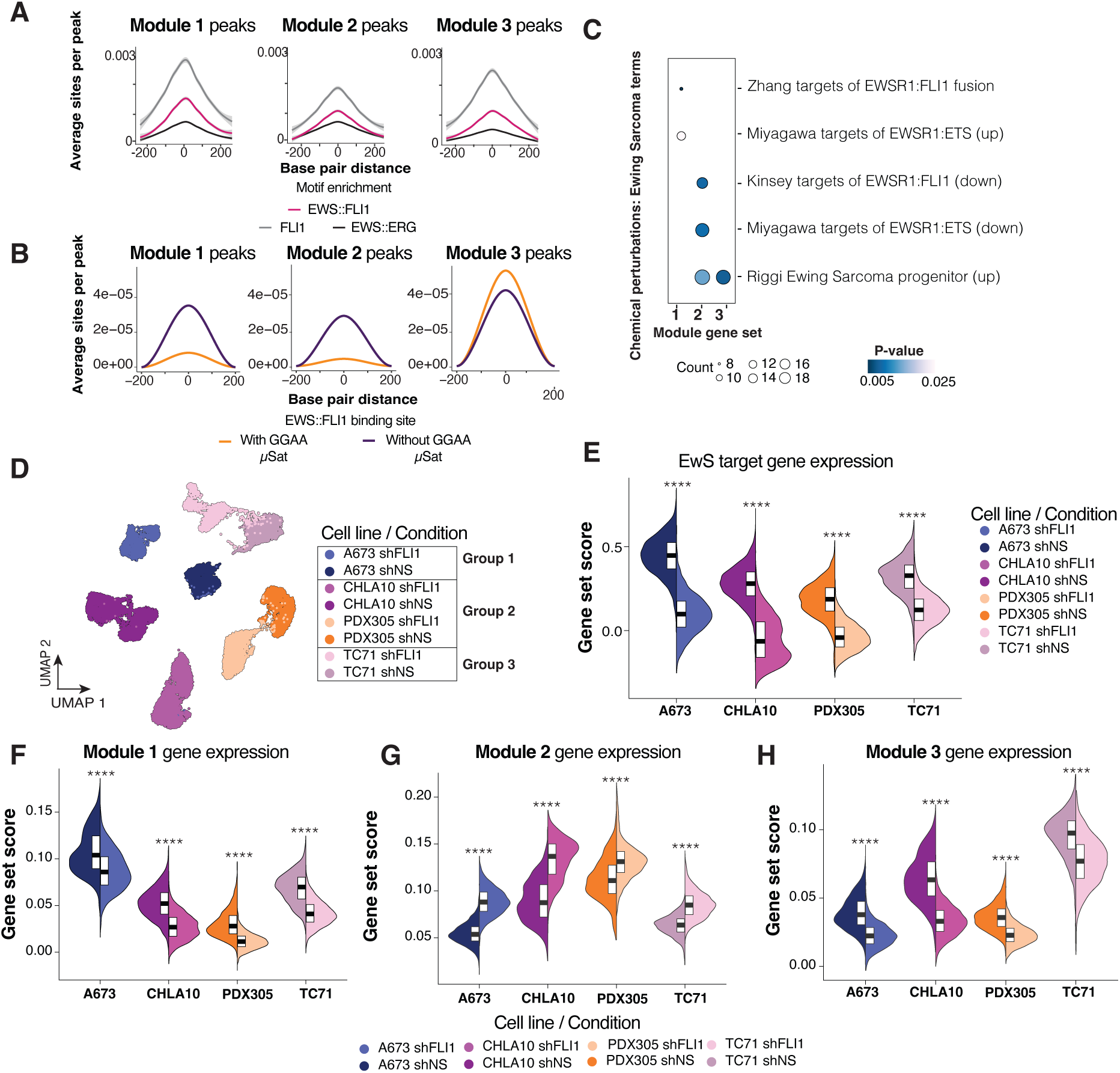
EwS gene regulatory modules are differentially regulated by EWS::FLI1. **(A)** HOMER enrichment of EWS/ETS motifs (EWS::FLI1, pink; FLI::ETS, gray; EWS::ERG, black) in module peak sets. **(B)** HOMER enrichment of EWS::FLI1 binding sites, separated by regions with (orange) or without (dark purple) GGAA microsatellites. **(C)** Universal enrichment analysis of MSigDB EWS::FLI1 and Ewing sarcoma terms in module gene sets. **(D)** UMAP embeddings of shNS and shFLI1 samples. EwS subgroup identity of cell lines represented by dashed boxes. Group 1 (A673) labelled with navy/dashed lines, group 2 (CHLA10, PDX305) in orange and group 3 (TC71) in light blue. **(E)** Expression of EwS target gene expression across shNS vs. shFLI1 samples. **(F-H)** Module gene set expression across knockdown samples. ****indicates p < 1×10-4 computed by a Wilcoxon Rank Sum test.

To begin to address the impact of EWS::FLI1 activity on cis-regulatory modules, we sourced data from published lists of EWS::FLI1 “activated” and “repressed” genes and performed gene set enrichment^33^ (Fig. 3C). Overall, EWS::FLI activated genes were enriched in all modules, whereas repressed genes were enriched only in module 2 (Fig. 3C). When combined with the lack of GGAA-uSat enrichment in module 2 (Fig. 3B), these observations are consistent with previous reports that repressed loci usually do not have long stretches of GGAA repeats^7,45^. Altogether, these data reveal distinct and polarized fusion-mediated regulatory events across EwS cell lines that differentially depend on GGAA repeats, suggesting that the diversity of fusion function is dependent on the motif landscape at fusion binding sites.

To directly address how different gene modules are affected by EWS::FLI1, we performed single-cell multiomic sequencing on selected cell lines from each EwS subgroup after shRNA- mediated knockdown of the fusion (Fig. 3D). Following quality control (Fig. S2A), we recovered 11,614 cells expressing negative control shRNA (A673 shNS, CHLA10 shNS, PDX305 shNS, TC71 shNS) and 12,466 cells expressing shRNA specific to the FLI1 transcript (A673 shFLI1, CHLA10 shFLI1, PDX305 shFLI1, TC71 shFLI1). All shFLI1 cells showed decreased expression of EwS TGs compared to shNS cells (Fig. 3E), providing evidence that the knockdown effectively induced loss of EWS::FLI1 activity. EWS::FLI1 knockdown had specific, predictable, and reproducible effects on all cell lines. Importantly the same stratification of shNS cell lines into groups with specific module enrichment was observed, indicating that lentiviral transduction and selection did not alter dominant gene programs (Fig. 3F-H, Fig. S2B-D). In addition, each module was affected by EWS::FLI1 knockdown in a manner consistent with predictions based on module gene sets. Specifically, EWS::FLI1 knockdown resulted in decreased expression of module 1 and 3 genes (Fig. 3F,H, Fig. S2B,D) and increased expression of module 2 genes (Fig. 3G, Fig. S2C). These data confirm that EWS::FLI1 activates cis-regulatory networks found in modules 1 and 3 but represses module 2. The results also support our initial hypothesis that EWS::FLI1-regulated gene modules differ among commonly used EwS cell lines. They further show that specific and reproducible patterns of transcriptomic and phenotypic signatures dominate each cell line and these predictably respond to fusion knockdown.

### CHLA10 exhibits intra-tumoral heterogeneity of gene regulatory modules

Most EwS cell lines in this study are dominated by a single gene-regulatory program (modules 1, 2, or 3). A notable exception is CHLA10, which contains mutually exclusive sub-programs defined by differences in both EWS::FLI1 motif activity and module utilization (Fig. 1B, Fig. 2A). This distinction is particularly striking because CHLA10 and CHLA9 originated from tumors from the same patient, with CHLA10 being derived post-therapy and following disease relapse. To further investigate this intratumoral heterogeneity, we performed unsupervised Louvain clustering^46^ on EwS cell lines (Fig. S3A) and computed the Pearson correlation coefficient of the most highly variable regions of accessibility. As expected, CHLA9, TC32, and PDX305 cells grouped together based on similarity (Fig. S3B). In contrast, CHLA10 segregated into two clusters (C22 & C23) that each showed greater transcriptional similarity to other cell lines than to each another (Fig 4A, Fig. S3B), highlighting the extent of its internal heterogeneity. Unsurprisingly, of all cell lines, CHLA10 C23 cells exhibited the greatest correlation to profiles from CHLA9.

**Fig. 4.**
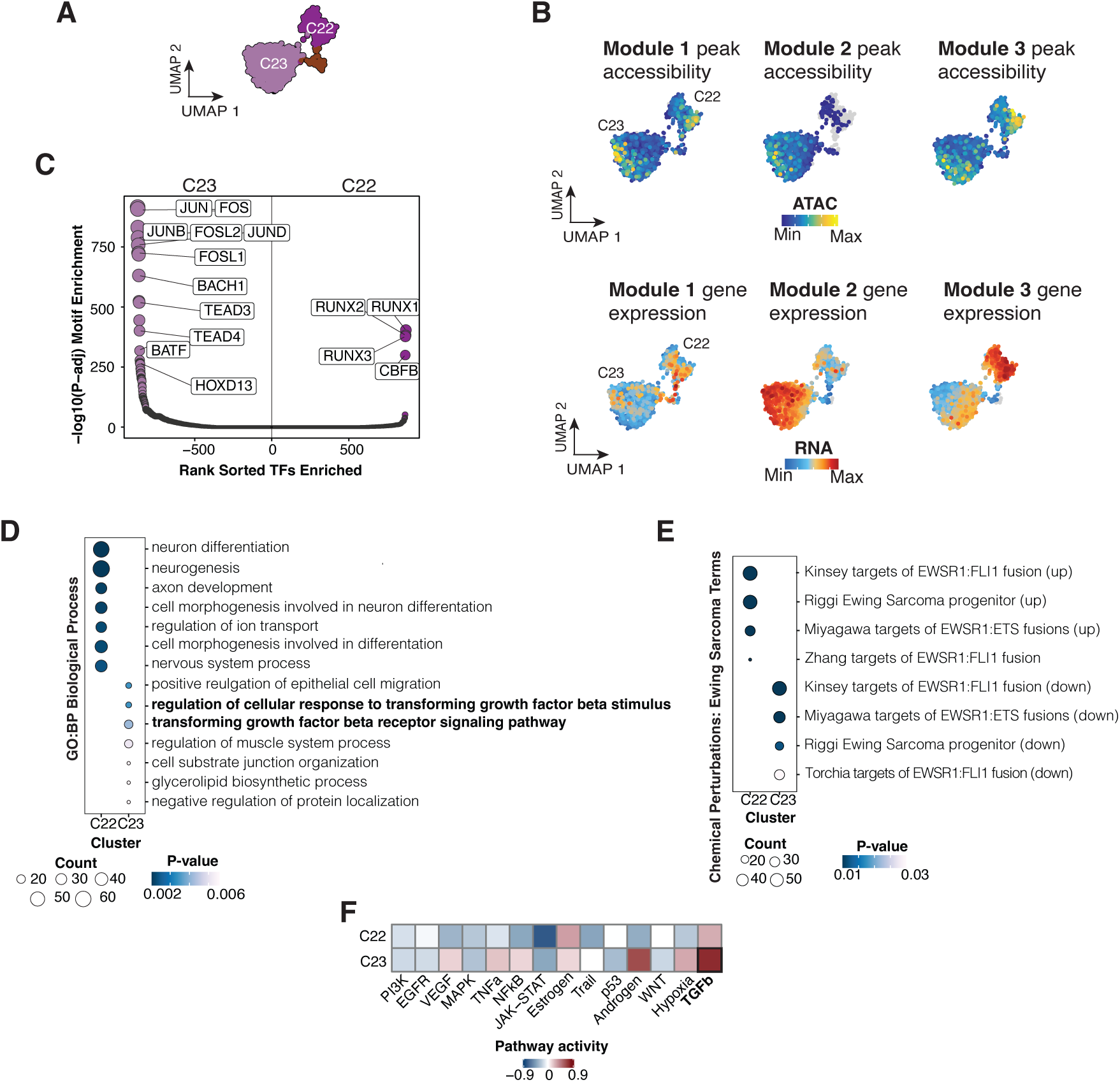
CHLA10 is a microcosm of EWS::FLI1 mediated heterogeneity. **(A)** UMAP of CHLA10 colored by cluster, C22 dark purple, C23 light purple. (B) UMAP of module peak accessibility and gene set expression in CHLA10 cells. (C) Differential motif enrichment between C22 and C23, TFs with -log10(p-value) > 250 highlighted. (D) Universal enrichment of GO: Biological Process terms in differentially expressed genes between C22 and C23, terms associated with TGF-b in bold. (E) Universal enrichment of MSigDB Ewing Sarcoma terms in differentially expressed genes between C22 and C23. (F) Pathway activity scores across C22 and C23, TGF-b highlighted in bold.

Cell-cycle effects contributed to but could not fully explain CHLA10 heterogeneity (Fig. S3C-D). Instead, plotting peak accessibility and transcript abundance of each module revealed that C22 cells enriched for module 3 while C23 cells relied on module 2 (Fig. 4B). Motif enrichment of differentially accessible peaks further underscored the nature of the split: RUNX motifs were enriched in C22 cells whereas AP-1, TEAD, and HOX TF motifs predominated in C23 cells (Fig. 4C). Gene-set enrichment mirrored these findings, C22 expressed neural programs whereas C23 cells were enriched for EMT, migration, and TGF-β signaling gene sets (Fig. 4D-E). Pathway-activity inference analysis between cell clusters^47^ further implicated TGF-β as a dominant upstream driver in C23 (Fig. 4F).

Collectively, the isogenic model CHLA9/CHLA10 encapsulates the spectrum of transcriptional states seen across the other EwS cell lines and pinpoints TGF-β signaling as key mediator of the module 2 transcriptional state.

### Sub-populations of EwS cell lines differentially respond to TGF-β stimulus

Prior reports have shown that TGF-β promotes pro-angiogenic extracellular-matrix (ECM) gene expression in EwS^48^. To interrogate the impact of TGF-β signaling on the newly defined transcriptional states in EwS, we performed multiomic profiling on A673 (module 1-dominant), CHLA10 (module 2 dominant), and TC71 (module 3 dominant) after treatment with recombinant TGF-β1 for 24 hours and compared them to vehicle-treated controls. After quality control filtering 19,611 cells were retained for analysis (Fig. S4A). Dimensionality reduction revealed substantial shifts in both gene expression and chromatin accessibility in A673 and CHLA10 following TGF- β treatment, whereas TC71 profiles remained relatively stable (Fig. 5A-C). In A673 and CHLA10, TGF-β treatment resulted in a clear polarization of cells towards module 2 usage (Fig. 5A-C), while expression patterns of modules 1 and 3 remained consistent with prior observations (Fig. S4B). The muted response of the TC71 cell line (module-3 dominant) to the influence of TGF-β (Fig. 5A-C, Fig. S4B) suggests that pre-existing transcriptional states may condition the sensitivity to TGF-β signaling.

**Fig. 5.**
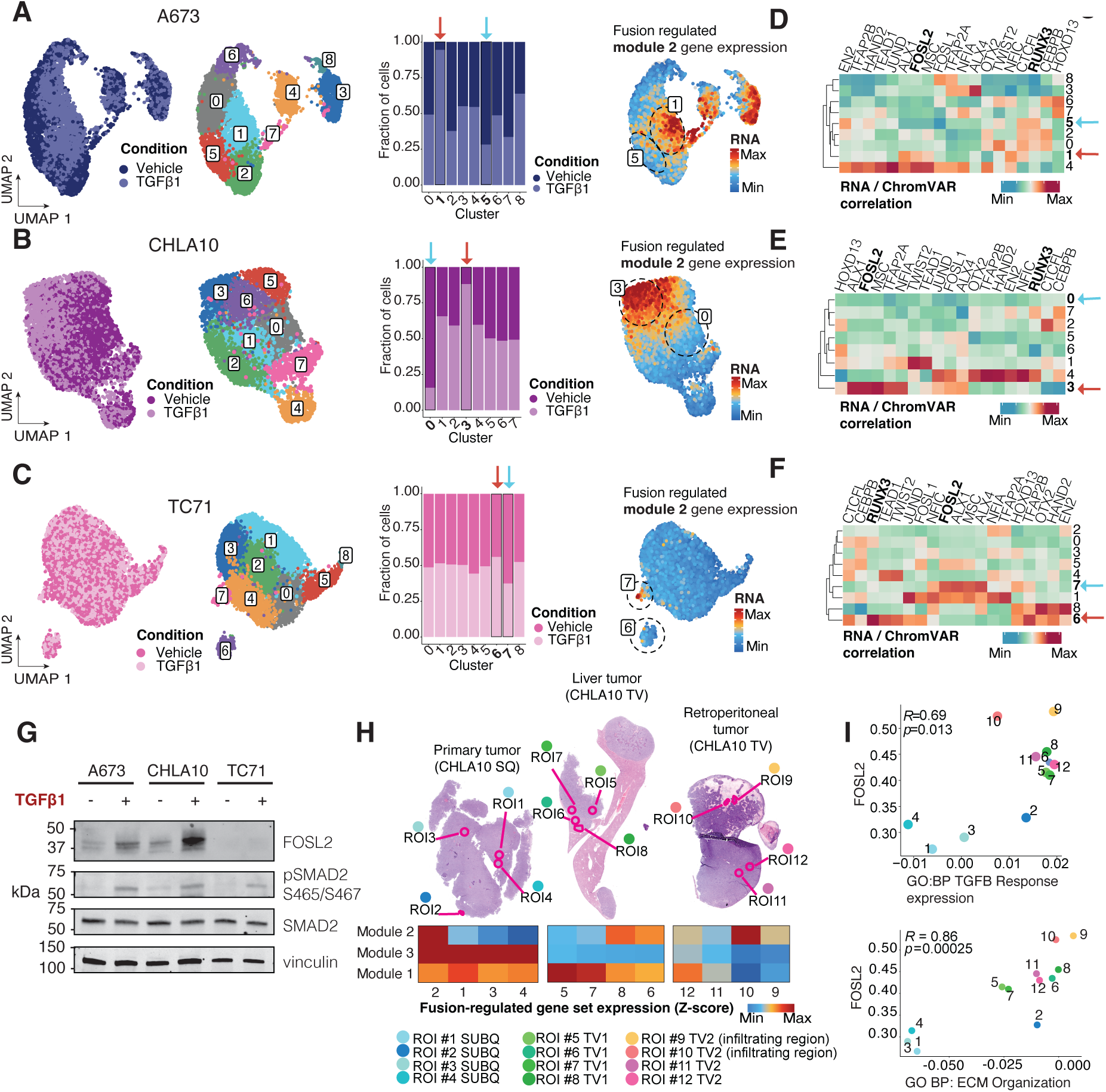
Sub-populations of EwS cell lines differentially respond to TGF-b stimulus. **(A-C)** UMAP of A673, CHLA10, and TC71 cells colored by condition (vehicle control or TGFb1 treatment) – *column 1*; and cluster – *column 2*; Fraction of treated versus control cells in each cluster with highest and lowest indicated by arrow - *column 3*; Expression of fusion regulated module 2 gene set, highest and lowest TGF-b clusters highlighted with dashed lines – *column 4*. **(D-F)** Correlation between TF expression and chromVAR TF deviation scores of EwS TFs across clusters, highest (red) and lowest (blue) TGF-b clusters highlighted by arrows. **(G)** Western blot of active (pS465/467) and total SMAD2 and FOSL2 protein in A673, CHLA10 and TC71 cells after 24 hr TGFb1 treatment. **(H)** H&E staining of 3 CHLA10 xenografts in NSG mice generated by subcutaneous or tail vein injection (*upper left*). Those regions of interest (ROIs) subjected to whole human transcriptome digital spatial profiling are highlighted. Heatmap of module gene expression across ROIs (*bottom left*). **(I)** Scatter plot of average expression of Gene Ontology: Biological Process (GO:BP) TGF-b response genes and extracellular matrix (ECM) organization genes compared to *FOSL2* expression in ROIs. Pearson R and p-value highlighted (*top left*).

We next examined TF expression and motif accessibility in clusters exhibiting the most pronounced or lowest enrichment of TGF-β-treated cells (indicated by arrows in Fig. 5A-F). Correlation analyses revealed a reciprocal enrichment pattern of many key TFs from Figure 2, where correlation of FOSL2 (but not RUNX3) gene expression and accessibility was generally highest in TGF-β responsive clusters. Western blot validation confirmed TGF-β-induced FOSL2 expression specifically in CHLA10 but not in the relatively insensitive TC71 cells, despite observed SMAD2 phosphorylation (Fig. 5G).

Our recent whole human transcriptome digital spatial profiling (DSP) study of CHLA10 tumor xenografts (Nanostring GeoMx) showed that transcriptionally distinct tumor cell subpopulations are spatially distributed^21^. We therefore interrogated these data to determine if the identified gene regulatory modules in CHLA10 cells were randomly or spatially distributed across tumors in vivo. We first selected genes from modules 1 & 3 that were significantly upregulated (average log_2_ fold change > 0.25) by EWS::FLI1 as well as module 2 genes that were significantly repressed by the fusion (average log_2_ fold change < -0.25) and applied these fusion-regulated gene sets to the DSP data. Intriguingly, regions of interest (ROIs) that were captured from tumor borders (ROIs 2, 6, 8 & 10) were enriched with tumor cells that displayed heightened module-2 expression, whereas densely packed tumor cores (ROIs 1, 3, 4, 5, 7, 11) favored modules 1 and 3 (Fig. 5H, Fig. S4C). Module 3 was also highly favored in subcutaneous compared to tail vein-derived disseminated tumors (Fig. 5H). Cells captured specifically from the border of a metastatic retroperitoneal tumor (ROIs 9 & 10) expressed the highest levels of *FOSL2*, TGF-β-response genes, and ECM-organization programs (Fig. 5I).

In summary, these findings highlight TGF-β-driven activation of module 2 gene expression in EwS cells via AP-1 transcription factors, particularly FOSL2. Furthermore, spatial analysis suggests module 2 programs are enriched at invasive tumor regions, underscoring the role of TGF- β signaling in mediating EwS cell heterogeneity and tumor progression dynamics.

### Patient tumors show heterogeneous usage of EwS cis gene regulatory modules

We next sought to determine whether the gene-regulatory networks defined in our cell line panel are similarly deployed across the landscape of early passage patient-derived-xenograft (PDX) and primary patient biopsies. To determine how the three regulatory gene programs manifest in these tumors, we used the fusion-regulated gene sets as defined above. In a series of publicly available PDX tumor profiles^19^, module 1 was prominent, while modules 2 and 3 (Fig. 6A) showed more modest and heterogeneous enrichment. On the other hand, in six primary EwS tumor samples^49^ (Fig. 6B, Fig. S4D), profiled by single-nucleus RNA-seq, every case contained discrete subpopulations that preferentially expressed either module 2 or module 3. In contrast to the PDX data, module 1-positive cells were comparatively rare in primary tumors.

**Fig. 6.**
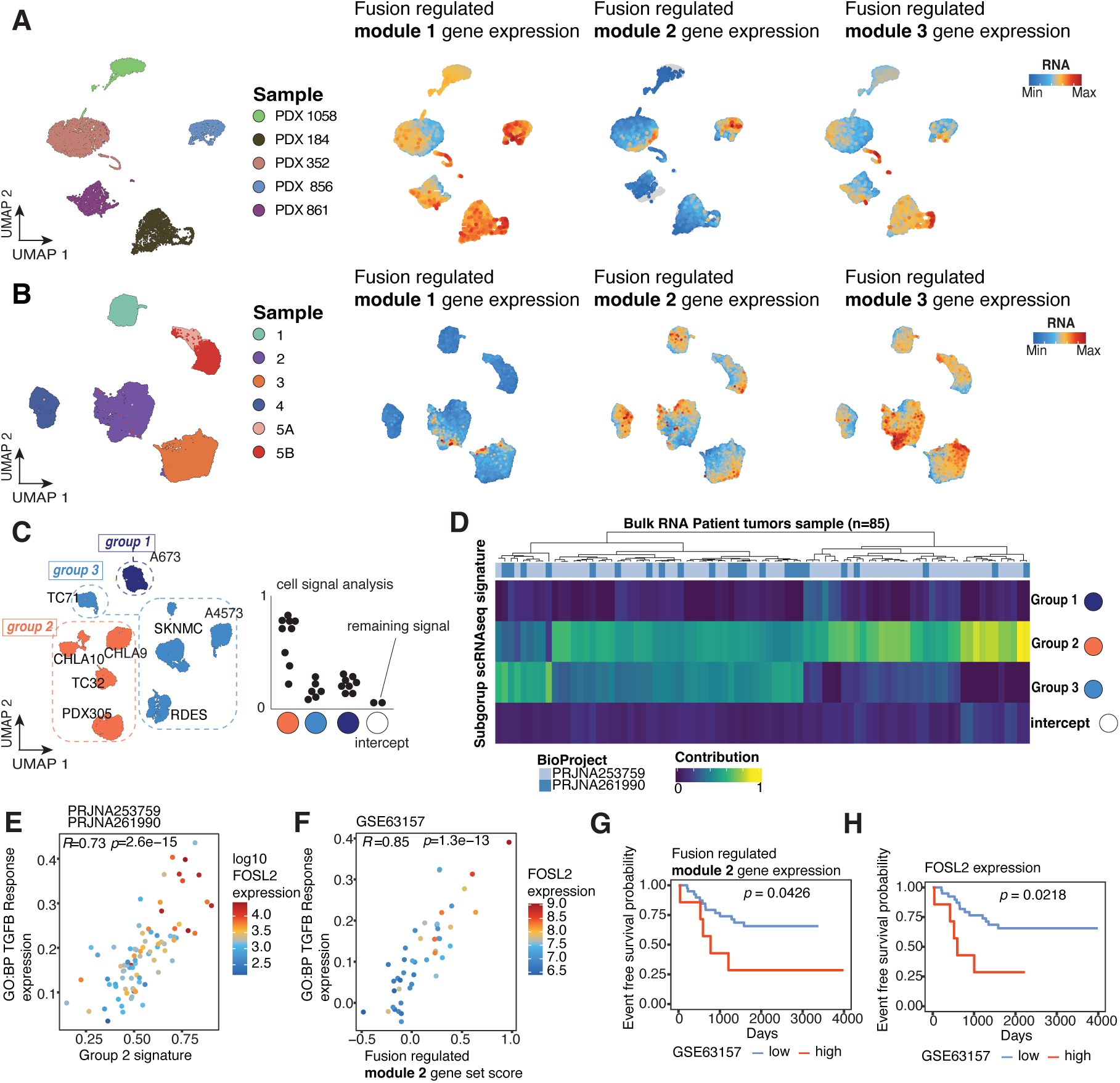
Human tumor data show usage of EWS::FLI1 gene regulatory modules. **(A)** UMAP embedding of five EwS PDX samples (*left*) UMAP of fusion regulated module gene sets in PDX samples (*right*) (B) UMAP embedding of single nuclei sequencing of 6 patient tumor samples (left). Patient tumor scRNA expression of fusion regulated modules (*right*). (C) Schematic depicting deconvolution method (D) Heatmap EwS subgroup signal in bulk EwS patient tumor RNA-seq. The intercept row represents remaining signal (bottom). Patient tumor samples annotated by GEO BioProject. (E) Scatter plot of bulk RNA patient samples by group 2 signature and GO: BP TGF-b response genes and colored log10 FOSL2 expression. Pearson R and p-value highlighted (*top left)*. (F) Scatter plot of microarray patient samples by module 2 expression and GO: BP TGF-b response genes and colored FOSL2 expression. Pearson R and pvalue highlighted (*top left)*. (G) Survival curves of patient microarray data51 (n = 46), high expression of module 2 gene set in red (top 15% expressing samples, n = 7), low expression in blue (bottom 85%, n = 39) (H) Survival curves of high expression of *FOSL2* samples (top 15%, n = 7, bottom 85%, n=39).

We next asked whether these single-cell patterns could be detected in bulk transcriptome data that were generated from three independent patient datasets^9,50,51^. To achieve this we employed a deconvolution method^52^ that identifies the overlap of signals from bulk measurements with a single-cell transcriptomic reference and captures any proportion of the bulk signal not represented by single-cell data in an ‘intercept’ term (schematized in Fig. 6C). We first validated the method on bulk RNA sequencing data taken from many of the same EwS cell lines we profiled in our experiments and found a high level of concordance with little signal captured in the intercept term (Fig. S4E). We then tested the transcriptomic overlap of EwS subgroups in 85 non-clinically annotated patient tumors that were profiled by bulk RNA sequencing in two independent cohorts (PRJNA253759, PRJNA261990). Group 2 signals were evident in nearly all tumors and group 3 cell signals were also present in more than half of tumors, confirming heterogeneity of gene network usage and cell state in primary tumors (Fig. 6D). Tumors with abundant group 2 signal also displayed significantly higher expression of TGF-β response genes and FOSL2 expression (Fig. 6E), linking our in-vitro observation that TGF-β drives module 2 activation to patient tumors in the clinical setting.

Finally, we examined microarray data from a series of 46 clinically annotated primary tumors (GSE63157) that were uniformly treated and for whom outcome data were available^51^. First, we observed significant correlation between expression of FOSL2, TGF-β response and module 2 expression (Fig. 6F). Furthermore, high expression of fusion-repressed module 2, and particularly of its key effector *FOSL2*, identified patients with markedly worse event-free survival, whereas modules 1 and 3 had little prognostic value (Fig. 6G-H, Fig. S4F). Module 2, linked to TGF-β responsiveness and FOSL2 activity, emerges as the predominant—and clinically adverse— program in primary EwS, underscoring its potential as both a biomarker of aggressive disease and a therapeutic target.

## Discussion

Single-cell genomics has made it possible to chart, with unprecedented resolution, the transcriptional circuitry that underlies intratumoral diversity^53,54^. In EwS, a cancer that has very few genetic mutations beyond its canonical chromosomal translocation^19–21,39^ and notable clinical heterogeneity, this diversity likely arises from epigenetic plasticity^55,56^. Our study enhances this understanding by defining two distinct transcriptional and chromatin accessibility states consistently observed across cell lines, patient-derived xenografts (PDXs), and primary tumor samples.

The first transcriptional state, associated with module 3 genes, is enriched for canonical GGAA-microsatellite enhancers, high EWS::FLI1 expression, and neural-crest transcription factors such as TFAP2A and RUNX3. In contrast, the second state associated with module 2 genes lacks GGAA repeats, is suppressed by EWS::FLI1, and involves activation by TEAD and AP-1 family members together with the homeobox factor HOXD13—all elements previously linked to epithelial-to-mesenchymal transition (EMT), invasion, and a low-fusion state^57–59^.

The mutually antagonistic architecture of these modules is consistent with earlier observations that EWS::FLI1-repressed loci tend to be short-repeat or single ETS sites that recruit additional repressor complexes such as ETV6^7,60,61^. Further supporting this, recent data^34^ demonstrate EWS::FLI1’s role in shaping cell identity by controlling promoter-enhancer looping and 3D chromatin networks. Loss of EWS::FLI1 was shown to alter 3D chromatin structure, activating enhancers enriched for AP-1 motifs and facilitating mesenchymal gene expression, consistent with our findings.

Analysis of three independent patient cohorts confirmed module 2 and module 3 signatures in most primary tumors. Clinically, elevated expression of module 2 genes, notably FOSL2, correlated with inferior event-free survival in a prospectively treated cohort of patients with localized EwS. This association nominates module 2 expression as a potential biomarker of aggressive disease and links TGF-β signaling to EMT-like transcriptional states and poor prognosis.

Another key insight from our study is the link between cells expressing high levels of module 2 genes and heightened responsiveness to TGF-β. These findings argue that external cues can re-wire specific subsets of EwS cells towards a more extreme mesenchymal/EMT-like state in the absence of genetic change. This is in line with emerging data that cell plasticity in EwS is influenced by a TGF-β-induced transition of EwS cells to cancer-associated fibroblast-like states^21^, which appear to be “unlocked” by TGFB2 response^62^. Thus, therapeutic strategies that disrupt these microenvironmental signals—or directly target downstream effectors such as FOSL2—may limit metastatic spread by constraining module 2 activation. Moreover, by promoting a fibroblast- like phenotype and dampening inflammatory signaling, TGF-β may also contribute to immune evasion and the establishment of an immunosuppressive tumor microenvironment^22,23,63^.

Our study is constrained by the limited number of primary single-cell datasets currently available; larger, clinically annotated cohorts will be required to validate the prognostic utility of module balance and to map temporal dynamics during therapy and relapse. Moreover, while CHLA9/CHLA10 serves as a useful isogenic model of plasticity, early-passage or organoid systems may preserve an even richer landscape of states. Finally, dissecting the full spectrum of extrinsic factors—beyond TGF-β—that tip the balance between modules will be key to developing microenvironment-targeted interventions.

In summary, our study provides novel insights into the TF regulatory networks that interact with the EWS::FLI1 fusion protein to sustain distinct cellular states and facilitate dynamic transitions between these states. Specifically, we identified core gene network modules and key TFs, notably FOSL2 and other module 2 TFs, which antagonize fusion-mediated repression, promoting a mesenchymal and EMT-like state associated with more aggressive disease phenotypes. Furthermore, our findings highlight cell-extrinsic TGF-β as a critical factor capable of activating FOSL2 and driving the aggressive module 2 program in vivo. By uncovering these regulatory mechanisms and their functional implications, we offer a foundation for developing therapeutic strategies aimed at limiting metastasis, overcoming treatment resistance, and ultimately improving clinical outcomes for patients with Ewing sarcoma.

## Materials and Methods

### Sex as a biological variable

Our study examined male and female animals, and similar findings are reported for both sexes.

### Cell Lines and PDX

The EwS cell lines A673, SKNMC, CHLA10, CHLA9, TC32, A4573, RDES, and TC71 were obtained from ATCC and COG (https://www.childrensoncologygroup.org/) cell lines repositories. The osteosarcoma cell line U2OS and rhabdomyosarcoma cell line RD were also obtained from ATCC. 5064L and 240L MSCs were obtained from Dr. Darwin Prockop. H7 MSCs were generously gifted from Dr. Alejandro Sweet-Cordero. PDX305 was generated from a recurrent metastatic tumor, as previously described^21^. A673, TC32, A4573, TC71, and SKNMC cell lines were maintained in RPMI 1640 media (Gibco) supplemented with 10% FBS (Atlas Biologicals) and 2 mmol/L-glutamine (Life Technologies). RDES was maintained in RPMI 1640 media supplemented with 15% FBS. Isogenic cell lines, CHLA10 and CHLA9, were maintained in IMDM media (Fisher) supplemented with 20% FBS, 2 mmol/L-glutamine, and 1X Insulin-Transferrin-Selenium (Gibco). PDX305 was maintained in RPMI 1640 media supplemented with 10% FBS, 2 mmol/L-glutamine, and 1% Antimycotic-Antibiotic (Gibco). U2OS was maintained in McCoy’s 5A media supplemented with 10% FBS and 2mmol/L-glutamine and RD was maintained in DMEM with 10% FBS. All hMSC lines were maintained in MEM-alpha media supplemented with 20% FBS, 2 mmol/L-glutamine, and 1% Antimycotic-Antibiotic. Cells were cultured at 37°C with 5% CO2. Cells were all confirmed to be mycoplasma free and identities subject to STR-confirmation every 6 months.

### Lentivirus Production and Genetic knockdown

pLKO.1 shNS (SHC002) and shFLI1 (TRCN0000005322) were used for FLI1 knockdown multiome experiments (Sigma). Knockdown plasmids were co-transfected with pCD/NL-BH*DDD (addgene, #17531) and pMD2.G (addgene, #12259) plasmids into 293FT packaging cells using PEI. 24 hours post-transfection, sodium butyrate was added to plates for 6 hours. Viral supernatant was collected 48 hours after transfection, filtered through 40 µM filters, and added the cells for 24-hours. Ewing sarcoma cells were selected by supplementing the media with 1.5 µg/mL puromycin (Gibco) for 48 hours before collection at 96 hours.

### TGF-β treatment prior to single cell sequencing

TC71, A673, and CHLA10 cells were plated at 500,000 viable cells per 10cm plate. 48 hours later recombinant human TGFβ1 (R&D 7754-BH-005, final dose 10 ng/mL in sterile 4 mM HCl and 0.1% BSA) or vehicle control was added. 24 hours later cells were trypsinized and >1×10^6^ viable cells were subjected to single cell library preparation and sequencing (details below).

### Western blots

Cells were washed with ice cold PBS, scraped and centrifuged, then lysed with RIPA buffer (Fisher Scientific) supplemented with protease and phosphatase inhibitors (Sigma) and sonicated (ten 30 second cycles). For TGF-β treated samples, recombinant TGFβ1 (10 ng/mL) or vehicle control (4 mM HCl, 0.1% BSA) were included in the media for either 30 minutes or 24hrs before lysis. Western blots were performed using the Bio-Rad Mini-PROTEAN Tetra System using 40 ug of lysate and 4-15% Mini-PROTEAN TGX polyacrylamide gels. Following transfer, nitrocellulose membranes were blocked in Odyssey Blocking Buffer (LI-COR) for 1 hour. Membranes were washed and incubated rotating overnight at 4°C with primary antibodies (FOSL2 CST #19967, HAND2 R&D AF3876, RUNX3 CST #13089, TEAD1 CST #12292, pSMAD2 S465/467 CST #3108, SMAD2 CST #3103, GAPDH Invitrogen AM4300, Vinculin CST #13901). Membranes were then washed four times in TBST for 5 minutes each and incubated with secondary antibodies (LI-COR IRDye 700CW or 800CW; 1:10,000) for 1 hour. After 4 additional 5 minute TBST washes membranes were scanned on a LI-COR Odyssey scanner.

### Digital Spatial Profiling

Q3 normalized gene counts from GeoMx Digital Spatial Profiling (NanoString) of EwS cell line xenograft tumors formed by CHLA10 injection were obtained from previous work^21^. This processed data was used to query fusion regulated module gene sets with respect to tumor microenvironment and injection site. Gene sets values for each ROI were calculated by converting the DSP Q3 counts matrix into a Seurat object and using the AddModuleScore function.

### Single-cell analysis

#### Cell and library prep

One million EwS cells were resuspended in PBS containing 2% BSA and nuclei were prepared according to manufacturer’s specifications. Cell lines in triplicate were pooled 1:1:1 at 1000 cells/uL and libraries were generated using the Chromium Next GEM Single- Cell Multiome ATAC+Gene Expression Kit following the manufacturer’s protocol (CG000338 Rev C). The resulting libraries were sequenced using a NovaSeq 6000 targeting a depth of 25K reads per cell (ATAC libraries) and 20K reads per cell (RNA libraries).

#### Single-cell multiome data analysis

Sequencing reads were demultiplexed and processed using the cellranger ARC pipeline. Cells with poor quality ATAC fragments and multiplets were filtered using ArchR’s standard workflow. Cells passing the following were kept for later analysis: percent mitochondrial RNA reads < 15%; 3 < log10(RNA/UMI counts) <= 4.5. Souporcell^64^ was used to identify cell lines and remove additional multiplets. Top RNA markers from scRNA data were compared to previously hashed CITEseq profiles^21^ to match genotype to cell line (GSE236289: <u>GSM7527501</u>, GSM7527502, GSM7527503). ArchR^37^, Signac^65^, and Seurat^46^ were used for downstream analysis. Dimensionality reduction was performed using addIterativeLSI() for both tile and gene expression matrices. An embedding was created by combineDims() and then addUMAP() with the mulitomic reduction. Group coverages and peaks calls were made using cell line identities. The motif matrix was built using addMotifDeviations(), addBdgPeaks(), and addDeviationsMatrix().

Average gene expression and gene activity scores were computed using addModuleScore(). EWS::FLI1 deviations were computed using the JASPAR2020 motif database. Transcription factor analyses used the cisBP database^66^. Highly utilized TFs were inferred using correlateMatrices(). All heatmaps were made with ComplexHeatmap. Marker peaks and genes were calculated using getMarkerFeatures() and getMarkersHeatmap(). Cis-regulatory elements were identified using addPeak2GeneLinks(). Universal gene set enrichment analysis was performed using clusterProfiler using the function enricher(), with MSigDB^67^ Hallmark and gene ontology references. Motif enrichment analysis on kmean peak data was performed using HOMER. The most enriched motifs were identified using findMotifsGenome.pl and motif enrichment was computed and visualized using annotatePeaks.pl.. EWS:ETS binding sites were sourced from previously published data^68^, and analyzed using HOMER and chromVAR with the same workflow as above.

Cells treated with TGF-β agonist and captured for multiomic sequencing were demultiplexed and processed using the same workflow as above. QC metrics used were identical to those above. A similar method of identifying each cell line using genetic demultiplexing was employed. Signac(*64*) was used to perform dimensionality reduction using weighted nearest neighbors. Motif matrices were constructed using RunChromVAR(), using cisBP^66^ as a database.

#### EWS::FLI1 long read sequencing

Starting material was 10x Genomics-generated full-length barcoded cDNA, leftover from what was used to create the Illumina-compatible libraries in the multiome downstream steps. Because 10x Genomics single-cell cDNA preparations typically contain a template switch oligo (TSO) priming artifact (i.e., TSO sequence at both ends of the cDNA molecule) instead of the correct structure, TSO artifact depletion was performed by using biotin-labeled primers for the streptavidin/biotin magnetic selection of molecules with the correct structure, as previously described^69^. To capture the gene transcripts encoding the EWS::FLI1 fusions, an enrichment was performed using xGen™ Hybridization and Wash v2 kit magnetic beads and reagents (IDT #10010351), as described in the manufacturer’s protocol, in addition to using 500 ng of cDNA starting material and using biotinylated capture probes designed with high specificity, tiling along exons 5, 6, and 7 of the *EWSR1* transcript, and exon 9 of the *FLI1* transcript. Following hybridization and magnetic capture of EWS::FLI1 molecules, an on-bead PCR was performed to amplify captured molecules, using standard 10x Genomic cDNA amplification protocol (described in 10x Genomics document #CG000315_Rev_B, Step 2.2) with the exception of a number of PCR cycles of 16 and an extension time at 72°C of 3 min. Following this, 500 ng of captured cDNA was DNA-repaired, A-tailed, SMRTbell adaptor-ligated, nuclease- treated, and SMRTbell-cleanup purified using the PacBio SMRTbell Prep kit 3.0 (PacBio #102- 141-700) as per manufacturer’s protocol. Libraries were sequenced on a PacBio Sequel IIe with 2 hours pre-extension time and 30 hours movie time.

Long-read computational processing pipeline consisted of retrieving error-corrected circular consensus sequence (CCS) BAM files generated by the on-board PacBio Sequel IIe (Pacific Biosciences of California, Inc.). CCS data were then processed by the programs in the IsoSeq3 suite of computational tools: 1) *Lima* to remove sequencing primers, 2) *Tag* to extract UMI and CB sequences, 3) *Refine* to remove polyA sequences, 4) *Correct* to error-correct CBs, and 5) *Dedup* to remove PCR duplicates, 4) *Pbmm2* to align reads. All the IsoSeq3 pipeline steps were used with default settings. Mapped BAMs were then used for pbfusion to detect fusions. Barcodes from long read molecules were extracted from mapped BAMs and merged with data of pbfusion.breakpoints.groups.bed files to count fusion transcript in single cells.

### External Data

The 78 direct EwS TGs were sourced from a previously published dataset^19^. Bulk RNAseq patient tumor samples were sourced from dbGaP under accession numbers PRJNA253759 and PRJNA261990. FASTQ files were aligned to reference genome (GRCh38) and using the piperna (https://github.com/furlan-lab/piperna) workflow which uses STAR^70^ aligner and generates a count matrix using the Bioconductor summarize overlap function (https://www.bioconductor.org) with its default settings. Microarray data was accessed from GSE63157^51^. RNA probes were annotated using pd.huex.1.0.st.v2, and normalized using oligo.

### scRNAseq Patient Tumors

Patient tumor biopsies were sourced and sequenced at St. Jude’s Children’s Hospital^49^. 10X reads were demultiplexed and processed using the cellranger pipeline. Ambient RNA was filtered out using cellbender. Cells with poor RNA measurements were filtered using the following thresholds: percent mitochondrial reads < 15% and 3 < log_10_(RNA counts) < 5.25. Dimensionality reduction and clustering was performed using standard workflow. Surrounding tissues cells were identified by using FindAllMarkers on Seurat clusters. Tumor enrichment was validated by choosing cells that express *CD99*, *NR0B1*, and EwS TGs. Identified tumor cells underwent normalization and dimensionality reduction with the same workflow. Average gene set scores were calculated using AddModuleScore().

### Deconvolution

Bulk RNAseq patient data were deconvoluted using cellSignalAnalysis (https://github.com/constantAmateur/cellSignalAnalysis). EwS subgroups 1-3 were used as transcriptomic references.

### Statistics

All statistical analyses were conducted using R or Python, with specific packages indicated in relevant sections of the Methods. Statistical significance was assessed using two-sided tests unless otherwise specified. For comparisons of gene set scores, motif deviations, or expression levels, Wilcoxon rank-sum or Student’s *t*-tests were used as appropriate based on data distribution and variance. Correction for multiple hypothesis testing was performed using the Benjamini-Hochberg method where applicable. Gene set enrichment analyses were conducted using the clusterProfiler package with MSigDB Hallmark and Gene Ontology annotations. For scRNA-seq and multiome analyses, differential expression and peak calling were performed using Seurat, ArchR, or Signac, and marker features were identified with adjusted *p*-values < 0.05 considered significant.

### Study approval

All work was approved by the Fred Hutchinson Cancer Center Institutional Review Board - Tumor Heterogeneity and Metastasis in Ewing Sarcoma - FHIRB0010627

## Data and materials availability

All code and processed data is available on https://github.com/furlan-lab/EwS_multiome. Data has also been deposited on GEO with accession GSE332630 including raw sequence. Single-cell RNA sequencing from primary samples are available at https://scpca.alexslemonade.org.

## Author contributions

Conceptualization: OGW, AA, EDW, JRS, ERL, SNF

Data curation: OGW, AA, EDW

Data generation: AA, EDW, MM, SB, SBK, AP, MD

Formal analysis: OGW, AA, EDW, RV, ZK, SNF

Software: OGW, RV, ZK, SNF

Visualization: OGW, AA, EDW

Supervision: JRS, ERL, SNF

Funding acquisition: ERL, SNF

Project administration: ERL, SNF

Writing—original draft: OGW, AA, EDW, AP, MD

Writing—review & editing: OGW, AA, EDW, RGG, RV, ZK, AP, MD, JRS, ERL, SNF

## Acknowledgments

The authors thank the Fred Hutchinson Scientific Computing Group and Genomics Core staff. This study was supported by multiple sources of funding, including the National Institutes of Health/National Cancer Institute (NIH/NCI) grant 1L40CA264534-01 (SNF), the 1Million 4 Anna Foundation (SNF, ERL), and NIH/NCI grant R01 CA215981 (ERL). Additional support was provided by NIH/NCI grant F31CA247104 (AAA) and the AACR-QuadW Foundation Sarcoma Research Fellowship in Memory of Willie Tichenor, Grant Number 22-40-37-WREN (EDW). Funding also came from the Sam Day Foundation (ERL), NIH grants S10-OD-020069 and S10-OD-028685, the Fred Hutch/University of Washington/Seattle Children’s Cancer Center Consortium NCI Cancer Center Support Grant P30 CA015704 and the Fred Hutch Translational Data Science Integrated Research Center Award (OGW).

## Competing interests

The authors declare that they have no competing interests.

## Notes

### Competing Interest Statement

The authors have declared no competing interest.

### Summary of Updates

Reviewers comments have been into version 3. Please see eLife VOR for full discussion

