## Supplemental Materials for "Multimodal single-cell analyses reveal distinct fusion-regulated transcriptional programs in Ewing sarcoma"

**Figure S1**

**A**

| Cell line/PDX name | Cancer/lineage type | age;sex | Fusion type | Tumor site |
|---|---|---|---|---|
| A673 | Ewing sarcoma | 15y;F | EWS::FLI1;type 1 | Muscle |
| A4573 | Ewing sarcoma | 17y;F | EWS::FLI1;type 3 | Pleural effusion |
| CHLA9 (isogenic with CHLA10) | Ewing sarcoma | 14y;F | EWS::FLI1;type 1 | Thoracic lymph node |
| CHLA10 (isogenic with CHLA9) | Ewing sarcoma | 14y;F | EWS::FLI1;type 1 | Thoracic lymph node |
| PDX305 | Ewing sarcoma | 17y;M | EWS::FLI1;type 1 | Abdominal soft-tissue metastasis |
| RDES | Ewing sarcoma | 19y;M | EWS::FLI1;type 2 | Humerus |
| SKNMC | Ewing sarcoma | 14y;F | EWS::FLI1;type 1 | Retroorbital metastasis |
| TC32 | Ewing sarcoma | 2.7y;F | EWS::FLI1;type 1 | Ileum and adjacent soft tissue |
| TC71 | Ewing sarcoma | 22y;M | EWS::FLI1;type 1 | Humerus |
| RD | rhabdomyosarcoma | 7w;F | N/A | Pelvic |
| U2OS | osteosarcoma | 15y;F | N/A | Tibia |
| 5046L | mesenchymal stem cell | 27y;M | N/A | Bone marrow aspirate |
| 240L | mesenchymal stem cell | 24y;M | N/A | Bone marrow aspirate |
| H7 | mesenchymal stem cell | 13y;M | N/A | Bone marrow aspirate |

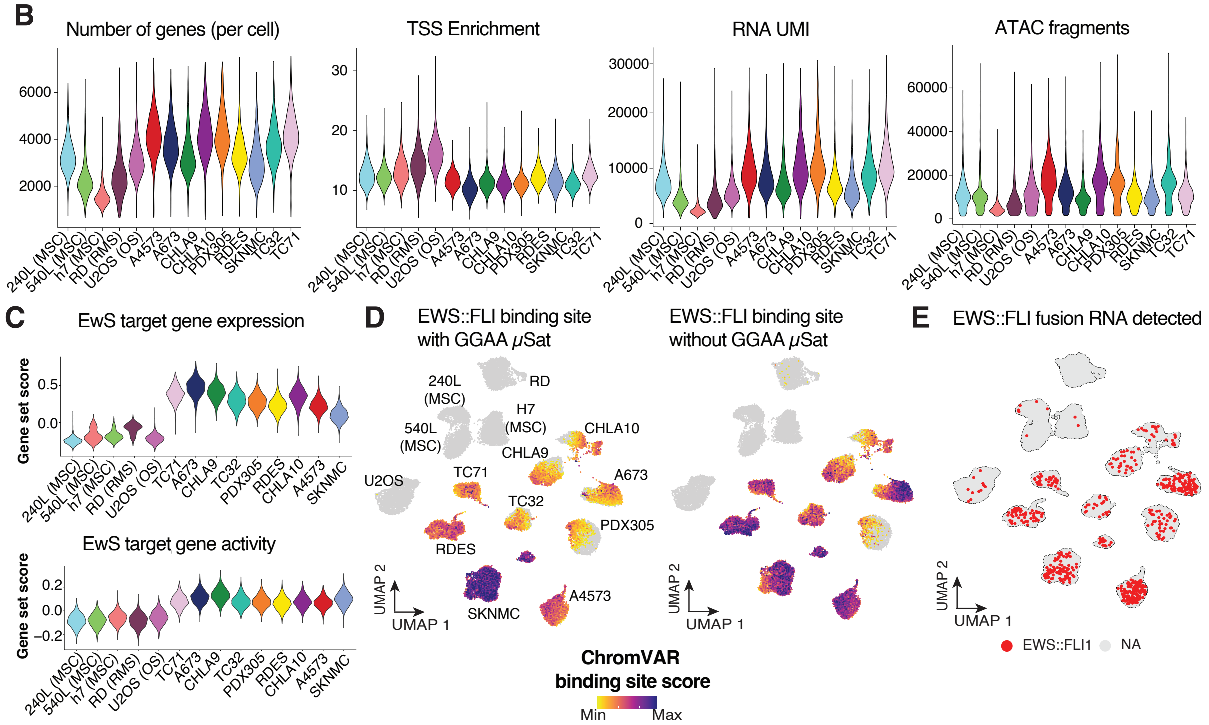


**Fig. S1. Multiomic analyses reveal distinct signatures between non-EwS and EwS cells. (A)** Table of all 14 cell lines used for the multiome sequencing. **(B)** Quality control metrics across cell lines. **(C)** Gene set expression and activity score of EwS target genes. **(D)** UMAP of all Ewing and Non-Ewing cell lines colored by chromVAR deviation scores of EWS::FLI1 binding sites, parsed by regions with or without GGAA microsatellites (μSAT; GGAA repeat ≥ 4). **(E)** UMAP of EwS and non-EwS cells highlighted (red) if EWS::FLI1 fusion transcript was detected by long-read sequencing.

**Figure S2**

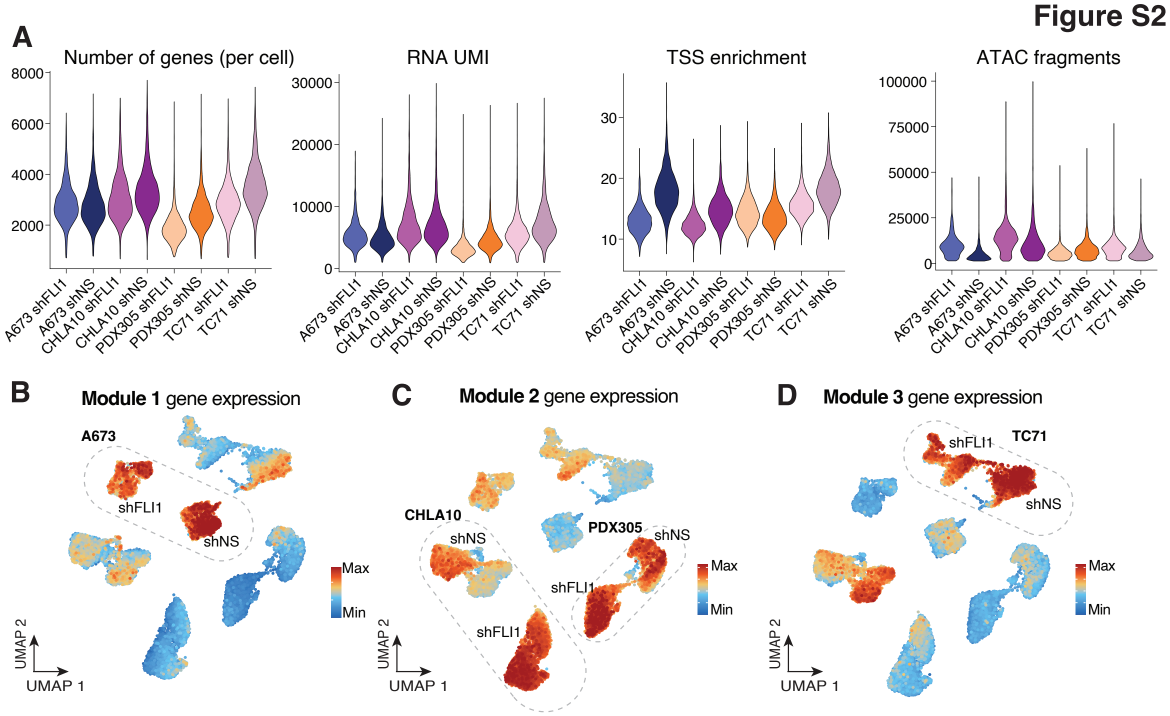


**Fig. S2. EwS gene regulatory modules are differentially regulated by EWS::FLI1. (A)** Quality control metrics of knockdown samples. **(B-D)** UMAP embedding of gene expression scores of module gene sets. Cell lines most enriched for a module gene set are highlighted.

Figure S3

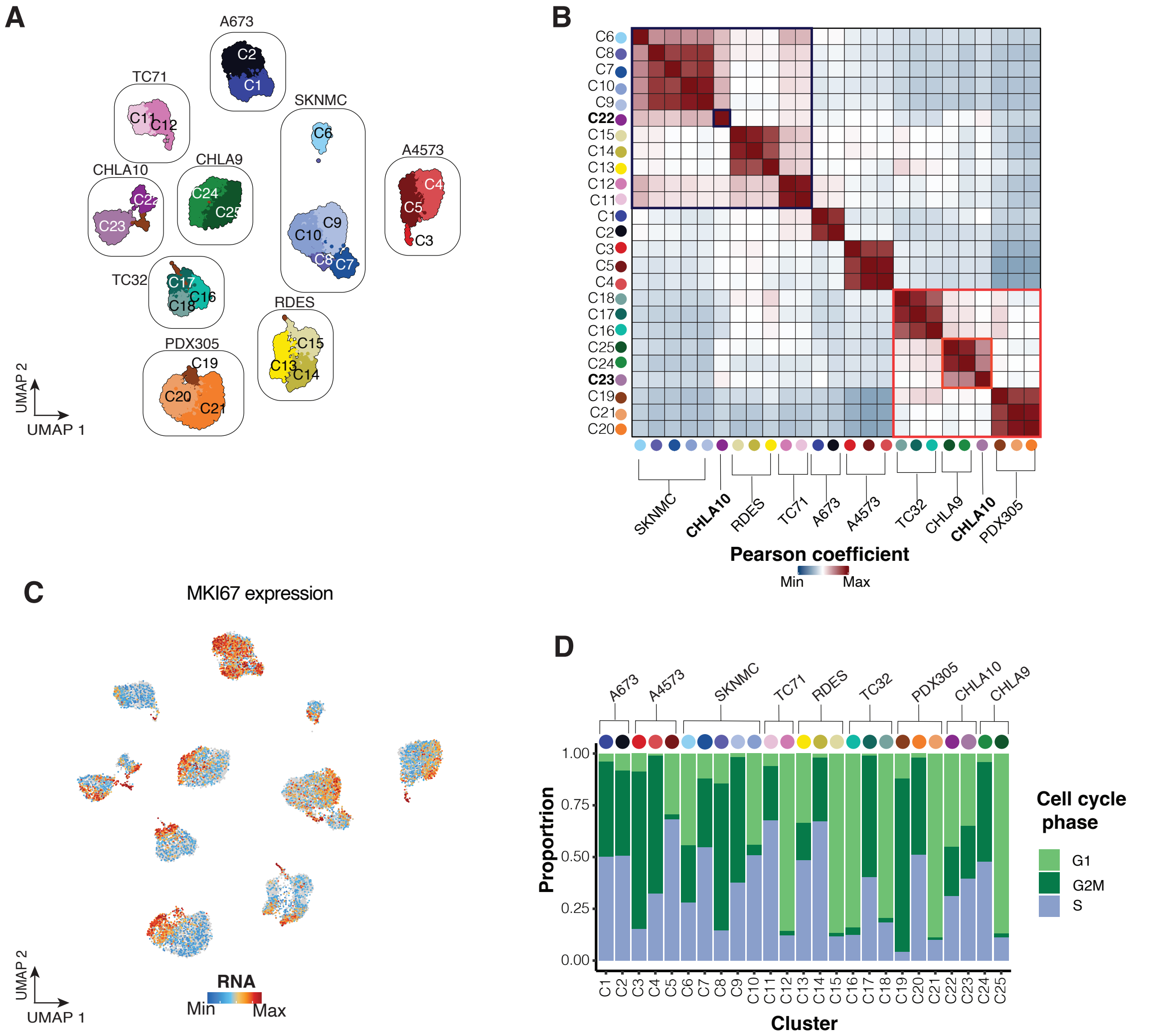


**Fig. S3. CHLA10 is a microcosm of EWS::FLI1 mediated heterogeneity**
**(A)** UMAP of 9 EwS cell lines colored and labeled by cluster **(B)** Pearson correlation heatmap of EwS clusters. The rows are labeled by cluster ID, the columns are labeled by cell line identity. Cell lines sharing epigenetic signatures highlighted in bold and red boxes. **(C)** *MKI67* expression across EwS cell lines. **(D)** Proportion of cells in G1, G2M, and S phase within EwS clusters.

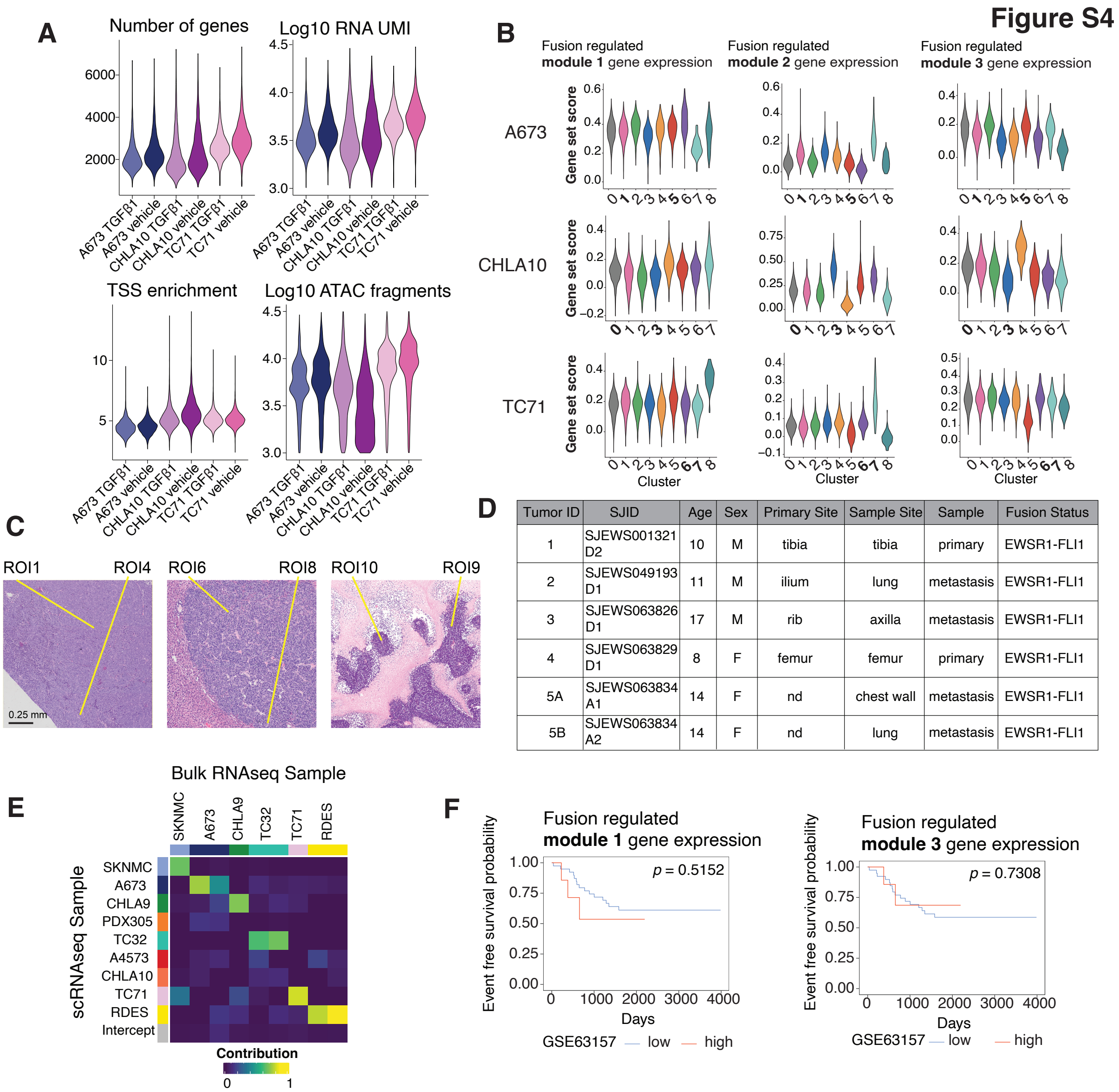


| Tumor ID | SJID | Age | Sex | Primary Site | Sample Site | Sample | Fusion Status |
|---|---|---|---|---|---|---|---|
| 1 | SJEWS001321 D2 | 10 | M | tibia | tibia | primary | EWSR1-FLI1 |
| 2 | SJEWS049193 D1 | 11 | M | ilium | lung | metastasis | EWSR1-FLI1 |
| 3 | SJEWS063826 D1 | 17 | M | rib | axilla | metastasis | EWSR1-FLI1 |
| 4 | SJEWS063829 D1 | 8 | F | femur | femur | primary | EWSR1-FLI1 |
| 5A | SJEWS063834 A1 | 14 | F | nd | chest wall | metastasis | EWSR1-FLI1 |
| 5B | SJEWS063834 A2 | 14 | F | nd | lung | metastasis | EWSR1-FLI1 |

**Fig. S4. Sub-populations of EwS cell lines differentially respond to TGF-β stimulus**
**(A)** Quality control metrics of vehicle control and TGF-β treated cell lines. **(B)** UMAP colored by gene expression of fusion regulated modules 1 & 3 across cell lines. **(C)** Zoomed-in imaged of ROIs 1, 4, 6, 8, 9 and 10 **(D)** Clinical details of patient samples subjected to single-cell sequencing. **(E)** Heatmap depicting similarity scores of publicly available bulk EwS cell line data with multiome scRNA-seq data. **(F)** Survival curves of patient microarray data(*1*) (n = 46), high expression of module genes 1 and 3 set in red (top 15% expressing samples, n = 7), low expression in blue (bottom 85%, n = 39)
